# Targeting CD19 for the treatment of autoimmune with a novel T cell engager

**DOI:** 10.1101/2024.02.17.580750

**Authors:** Andy Qingan Yuan, Likun Zhao, Lili Bai, Yang Wang, Isabella Yang, Ray Qirui Yuan, Qingwu Meng, Xuning Dai

## Abstract

Autoimmune disorder, affecting more than twenty million in United States and billions of people worldwide, occurs when human immune system mistakenly attacks its own body. Autoreactive B cells, particularly the autoantibodies they produce, play a critical role in the etiology, progress, prognosis, and pathogenesis of multiple autoimmune diseases. While targeting B cells medicines, such as anti-CD19 or anti-CD20 monoclonal antibodies have produced promise in both pre-clinic models and some clinic practices, many patients failed to respond, ended less satisfactory outcome, or relapsed to monoclonal antibody therapies. Novel therapies are desperately needed to fight this challenge. In this study, based upon a symmetrical format, namely fusion of IgG and scFv technology (FIST), an anti-CD19xCD3 T cell engager (TCE) YK012, developed to treat B-cell malignancies such as precursor pre-B acute lymphatic leukemia (ALL) and non-Hodgkin’s lymphoma (NHL), was expanded to explore the treatment of autoimmune diseases by bringing T cells to CD19 positive B cells, including autoreactive B cell populations. Even as bivalent binding to CD3, YK012 has so significantly reduced binding affinity to TCR compared to other TCEs that, free YK012 alone does activate T cells at all. In contrast, in the presence of CD19-expressing Raji target cells and mediated by YK012, T cells can be activated in a dose-dependent manner, measured by upregulation of NFAT-reporter, CD25, IFN-γ and granzyme B expression. Crosslinking of B cell with T cell by YK012, the conjugation elicited T cell-mediated potent but mild cytotoxicity only to CD19-positive Raji cells. Additionally, YK012 elicited much reduced cytokines release *in vitro* compared to that of Blincyto^®^ biosimilar, implying a potentially me-better therapeutics replacement in clinics for B-cell related diseases. In two *in vivo* pharmacological studies, YK012 alleviated the pathology symptoms of a collagen-induced arthritis model and almost completely depleted the peripheral B cell in human stem cell-transplanted murine model. In summary, this study demonstrated that YK012, a FIST platform designed TCE, has the potential to treat B cell-mediated diseases including autoimmune indications with possible better efficacy and less side effects in clinical practices.

## Introduction

An autoimmune disorder occurs when a person’s immune system mistakenly attacks its own body due to the failure of self-tolerance^1^. There are more than 80 types of autoimmune diseases, which affect 23.5 million Americans and have devastating impacts on health, well-being, and quality of life. In normal conditions, B cells are critical components of adaptive immune system, besides the assist roles in promotion of immune reactions, mainly responsible for generating antibodies, which help to eliminate the pathogens^2^. When self-tolerance barrier is broken, however, auto-antigens activated reactive B cells often produce a large amount of autoantibodies, which play a key role in the pathogenesis of a wide variety of autoimmune diseases, such as systemic lupus erythematosus (SLE), rheumatoid arthritis (RA), multiple sclerosis (MS), and neuromyelitis optica (NMO), membranous nephritis (MN), to name a few.

The pathogenic roles for B cells vary across autoimmune diseases, which can be autoantibody dependent or independent^1, 3^. Nevertheless, reduction or total depletion of autoreactive B cells has shown promises for multiple autoimmune diseases in both preclinical models and clinical settings^3^. B cell targeted monoclonal antibodies have been approved for the treatment of RA, SLE, MS etc. by FDA, and the indications of autoimmune diseases that can be treated with such strategy are still expanding^3, 4^. One major successful approach to target B cells is to directly deplete B cells with monoclonal antibodies against CD20, such as rituximab and ocrelizumab to induce Fc-mediated ADCC. Despite its approval, the responses to rituximab vary in a wide range and a substantial proportion of patients do not show improvement of disease severity^3, 5, 6^. Rituximab can largely deplete B cells in the periphery blood stream; however, it is less efficient to eradicate the autoreactive B cells in the lymphatic organ and inflamed tissues^6^. The mechanisms underlying the impaired depletion by anti-CD20 in lymphoid and inflamed tissues are not yet well understood, probably because minimal CD20 density is required for ADC^7,8^. Indeed, CAR-T therapy targeting CD19-expressing B cells with CAR-modified T cells, working at extremely low antigen density^9^, has overcome the limitations of mAb-based therapy in refractory SLE patients in pilot trials^10^. The MOA of CAR-T is cell-bridging, i.e. the conjugation of T with B cells. In that sense, bispecific antibody-based T cell engagers targeting autoreactive B cells beyond CAR-T, being off-the-shelf and cost-efficient by nature, are better positioned to offer a more applicable approach for treatment of poor-responsive, relapsed, and refractory autoimmune patients.

While CD20 is expressed by mature B cells, CD19 is expressed throughout B cell development and maturation including mature B cells, and some pre-B cells, plasma blast cells, and plasma cells^11^. The polyreactive and thus autoreactive B cells are predominantly selective in the early B cell development stage, and they do not express CD20^3^. In addition, the long-lived plasma cells having high CD19 and low CD20 expression, are resistant to B-cell depletion therapy^3^. Overexpression of human CD19 in transgenic mice results in hyperactivation of B cells with elevated humoral response, disrupted tolerance and spontaneous induction of autoimmunity^12^. CD19 expression is 20% higher on B cells from some patients with MS compared with healthy controls and the levels of expression are correlated with disease susceptibility^13^. Thus, targeting B cells via CD19 with TCEs may allow broader and deeper elimination of B cells, including CD19^+^CD20^lo^ plasma blasts and CD19^+^ pre-B cells, which could provide better outcomes than anti-CD20 approach for autoimmune disease patients.

Bispecific T cell engager (BiTE) is a pioneering format of TCE molecules that simultaneously binds an antigen on a target cell and CD3 on a T-cell. BiTE brings T cells to the proximity of the B cells, resulting in activation of T cells that can subsequently kill the target cells^14^. The first-in-class BiTE, blinatumomab (Blincyto^®^), is a fusion protein composed of two single-chain antibodies (scFvs) and has been approved for the treatment of patients with relapsed and/or refractory B cell-precursor ALL^11^. Owing to its small size and strong affinity to CD3, blinatumomab has a very short half-life and strong toxicity^15^. So far, TCEs have been extensively studied for the treatment of various cancers and have yet to be investigated for the treatment of autoimmune diseases. We have designed and engineered a novel bispecific anti-CD3xCD19 TCE, named YK012, based-upon the in-house FIST (fusion of IgG and scFv technology) platform. YK012 has significantly weak binding to CD3 and reduced potency to activated T cells but maintaining the capabilities to bridge B with T cells and redirected killing of autoreactive CD19^+^ B cells. It induced low levels of cytokine release *in vitro*, potentially carrying a better safety profile than Blincyto^®^ so that besides its original intention for treatment of blood cancers, it might have suitable applications in autoimmune indications.

In this study, we characterized and investigated the potential of YK012 for the treatment of autoimmune diseases by killing the B cells in *in vitro* and *in vivo* models. *In vitro*, YK012 only induced activation of T cells in the presence of CD19^+^ Raji B cells and initiated T cell-mediated killing of CD19^+^ Raji B cells but not MV-4-11 CD19^-^ monocytic leukemia cells. YK012 elicited much reduced *in vitro* cytokines release compared to that of Blincyto^®^ biosimilar. In two *in vivo* pharmacological studies, YK012 alleviated the pathology symptoms of a collagen-induced arthritis model and almost completely depleted the peripheral B cell in human stem cell-transplanted murine model. In summary, the report validated the advantages of FIST platform in developing ideal novel TCEs, revealed the potential of YK012 for the treatment of autoimmune diseases, and provided foundation for further clinical trials of this candidate in various B-cell related indications.

## Materials & Methods

### Cells, animals, reagents and culture medium

Raji cells, Jurkat T cells were purchased from ATCC and cultured in RPMI Medium 1640 (Gibco, 21870) supplemented with 10% FBS, 2 mM L-glutamine, and 100 U/ml penicillin and streptomycin (Gibco, 15070063). Raji-luc and NALM6-luc were developed in house from Raji or NALM6 respectively with stable expression of luciferase in each. Raji-lp was derived in-house from Raji integrated with stable expression luciferase and puromycin. NFAT-Jurkat E6.1 was derived from Jurkat cells that have been transduced with a luciferase gene driven by the NFAT transcription factor. Human PBMC were purchased from Allcells Biotechnology (Shanghai) Co., Ltd. (PB003-C-10M). Human PBMC were cultured in X-VIVO^15^ medium (LONZA, 04-418Q).

Blincyto^®^ biosimilar was prepared in-house according to its official amino acid sequence. UCHT1 mAb was purchased from ThermoFisher Scientific (Product # 16-0038). Cytokine release detection kits, including human IFN-γ ELISA Kit (Cat# 1110002), human TNF-α ELISA Kit (Cat#1110202), human IL-2 ELISA Kit (Cat#1117202), human IL-6 ELISA Kit (Cat#111002) were purchased from Dakewe Biotech Co., Ltd. Transgenic mice hCD3EDG/hCD19 induced CIA model were provided by Suzhou Hekai. HuHSC-NCG-M mice were developed by Gempharmatech.

### YK012 design, expression, and purification

The variable sequence of the anti-CD19 and anti-CD3 were derived and humanized from patent-expired clones, HD37 and UCHT1, respectively. The tetravalent bispecific antibody YK012 was built by FIST platform, optimized in Excyte Biopharma Ltd. It is composed of an anti-CD19 IgG4P unit and two anti-CD3 scFv units, fused at the C-terminus of Fc through a flexible linker peptide. S228 to P mutation was introduced in the hinge region to promote the stability of final product. The YK012-coding gene was assembled as classical IgG in an in-house vector pG4HK by standard molecular cloning methods. The plasmid was stably transfected into CHO-K1 (Shanghai Biochemical Institute, China Academia of Sciences) suspension cells. High expression clones were selected, and fermentation supernatants were purified by Protein-A affinity chromatography using MabSelect™ column (GE Healthcare, Princeton, NJ). The bispecific antibody was further polished by cation exchange chromatography using Capto MMC (GE Healthcare, Princeton, NJ) and ion exchange chromatography using Q HP (GE Healthcare, Princeton, NJ) columns. The purified protein was concentrated and stored in 50mM PBS containing 0.15M NaCl buffer. The purity was analyzed with intact mass spectrometry, SDS-PAGE and size-exclusion chromatography (HPLC-SEC). Blinatumumab (Blincyto®) biosimilar was generated and prepared in mammalian cells in house according to its published amino-acid sequence (Blinatumomab: Uses, Interactions, Mechanism of Action | DrugBank Online). The YK012 drug substance was stored at -80 °C.

### Binding characterizations of YK012 to CD3 and CD19

The binding affinities of YK012 to human CD3 ε/δ heterodimer (ACROBiosystems, CDD-H82W6) and CD19 (Sinobiological, 11880-H08H) were measured by BIAcore. The cellular binding KDs of YK012 to CD3 on Jurkat T cells, human T cells, and CD19 on Raji-lp cells and human B cells were determined by standard flow cytometry or receptor saturation assay, respectively. In brief, the cells were washed twice with PBS and incubated with FACS buffer (1×PBS with 0.5% BSA) in a 96-well U bottom plate containing various concentrations of YK012 for 30 minutes at room temperature. Subsequently, the cells were washed twice with FACS buffer and stained with Alexa Fluor 488-conjugated goat-anti human IgG Fc antibody (Jackson Laboratory, 109-545-098). The binding of mouse anti-human CD3 (clone: UCHT1) to Jurkat T cells, Alexa Fluor 488-conjugated goat-anti mouse IgG Fc antibody (Jackson Laboratory, 115-545-003) was used for the second antibody. Samples were analyzed by flow cytometry with a Guava Technologies flow cytometer (Guava^®^ easyCyte 8HT). To measure the apparent binding affinity of YK012 to human Jurkat and Raji-lp cells, saturated method^22^ was applied with titrated YK012 added to the two cells, respectively.

### Peripheral blood mononuclear cell (PBMC) isolation and activation of T cells

PBMCs were isolated from healthy donor buffy coat and isolated by Ficoll density gradient centrifugation. PBMCs were cultured in complete RPMI 1640 medium supplemented with 10% heat-inactivated fetal calf serum (Gibco, 16000044), 100 U/ml penicillin and streptomycin (Gibco, 15070063), 1 mM pyruvate (Gibco, 11360070), and 55 µM β-mercaptoethanol (Sigma, m3148). Cells were activated by TCR stimulation using plate-bound human anti-CD3 and soluble CD28 (clone OKT-3 and clone 9.3 Bio X Cell, respectively, 5 µg/mL of each) for 3 days, followed by interleukin 2 (IL-2) (20 ng/mL, Gibco) stimulation for 5 days.

### Determination of T cell activation by flow cytometry

For measurements of T cell activation and cytokine expression, T cells from human donors were cocultured with at an E:T ratio of 10:1 or without Raji cells in the presence of 1 µg/ml YK012 for 72 hours. T cell activation was assessed by incubating cells with fluorescein isothiocyanate-conjugated anti-human CD8 (CD8-FITC, Biolegend, 100705) and phycoerythrin-conjugated anti-human CD25 (CD25-PE, Biolegend, 356103).

For cytokine expression, cocultured cells were stimulated with PMA, ionomycin and GolgiPlug for 4 hours, and antibodies against phycoerythrin-conjugated anti-IFN-γ (Biolegend, 383303) and fluorescein isothiocyanate-conjugated anti-granzyme B (Biolegend, 372205) were used for intracellular staining. Samples were analyzed by flow cytometry with a Guava Technologies flow cytometer.

### T cell proliferation assay induced by YK012

Human peripheral blood mononuclear cells (PBMC) were isolated from healthy donors, purified T cells (panT) were also prepared. PBMC or panT Cells were labeled with CFSE (final concentration of 5 µM of in PBS) at room temperature for 7 minutes in the dark. The labeled cells were then washed with PBS containing 10% serum and seeded at 1×10^6^ cells/well into a 48-well plate. Different concentrations of YK012 (final concentrations of 100, 200, and 400 ng/ml) were added to each culture wells. After 120 hours of incubation, the division and proliferation of PBMC or T cells were examined using flow cytometry. The positive control was anti-CD3×CD28 activation beads (5 µl/ml), while the negative control was set by omission of stimulation reagents. To distinguish between the CD4 and CD8 subpopulations of T cells in PBMC or panT, the sample cells were divided into two fractions. One fraction was stained with CD4-PE to detect the division of CD4+ T cells, and the other fraction was stained with CD8-PE to assess the proliferation and division of CD8+ T cells.

### YK012 activates T cells through NFAT pathway

Jurkat E6-1-NFAT-luc cells are Jurkat cells that have been transduced with a luciferase gene driven by the NFAT transcription factor. After the activation of the NFAT signaling pathway, the reporter gene luciferase is expressed, and upon interaction with luciferase substrates, a detectable signal (RLU) is produced. The intensity of the fluorescent signal is related to the extent of T cell activation, making it a sensitive and simple method for detecting T cell activation.

Jurkat E6-1-NFAT-luc cells were adjusted to a density of 2×10^6^ cells/ml, and Raji cells were adjusted to a density of 0.2×10^6^ cells/ml. 50 µl of each cell suspension was added to wells of a 96-well plate. 100 µl of YK012, started at a concentration of 4 nM and 4-fold diluted in X-VIVO^15^ medium was added to each well. The eleventh column was left without any antibody. The cells were cultured at 37°C with 5% CO_2_ for 24 hours, after which the assay was performed to detect the fluorescence signal.

### Cytokine release induced by YK012 or Blinatumomab, with or without dexamethasone

Liquid-nitrogen-stored human PBMC was immediately thawed at a 37℃-water bath and washed with PBS twice to remove storage medium. The cells were resuspended in X-VIVO to 2×10^6^ cells/ml and dispensed at 50 µl/well to a 96-well culture plate. Raji cells at 2×10^5^ cells/ml were added to the each well at 50 µl/well to reach PBMC: Raji =100000:10000 (10: 1). A blank control well without Raji was set. Stock YK012 or Blinatumomab biosimilar, adjusted to 8 nm, was diluted at 4-fold down and added to each well at 100 µl, in a duplicate manner. A constant 100nM dexamethasone was applied to each well for the effect on cytokine release in a separate plate. The plates were incubated at a cell culture incubator for 48 hours after gentle mixing. An ELISA method was applied to measure the concentrations of individual IL-2, TNF-α, IFN-γ, IL-6 released in each well. Plots of each cytokine level verse YK012 or Blinatumomab biosimilar concentration, with and without dexamethasone, were generated by Microsoft Excel.

### Cytotoxicity study of YK012 over time and on different cells lines

The healthy human PBMCs were activated with anti-human CD3×CD28 magnetic beads (Novoprotein Technology, Co., Ltd, GMP-B038). 1 µl magnetic beads were added for every 1×10^6^ cells. After 3 days of culture the magnetic beads were removed through a magnetic rack from the culture, which could be expanded more to produce enough activated human T cells. 10 ml (10^6^ /ml) human T cells were harvested and washed twice with PBS for each 96-well plate. The human T cells were adjusted to 2×10^6^ cells/ml with X-VIVO^15^ and aspirated to a 96-well U-bottom plate at 50 µl/ well. Raji-luc or NALM6-luc were constructed in-house to stably express luciferase gene. Target cells (Raji-luc or NALM6-luc) were prepared and adjusted to 1×10^6^ cells/ml. The cells were washed twice with PBS, resuspended to 0.2×10^6^ cells/ml, and added to a 96-well plate at 50 µl/well. Fixed E:T ratio (10:1) was used in the experiment.

Starting from 8nM, YK012 was diluted by 4 folds (1x volume of antibody were added with 3x volume of X-VIVO^15^ medium) to 10 grades and 100µl of each diluent was added to the target cell wells in duplicate mode. X-VIVO^15^ medium control (no antibody added or 0 nM concentration) was set as blank. The mixture was placed in a 37℃, 5% CO_2_ cell incubator, sampling at different culture time points (2, 4, 6, and 24 hours) for potency (EC_50_) calculation. Before the samples were taken for reading, cells were from each well evenly stirred with a 200µl-pipette and 60µl was transferred to opaque white reading plates. 60µl luciferase substrate was added to each well, mixed well, and read using Synergy HT fluorescence reader in fluorescence intensity mode to obtain the values (RLU).

Data analysis method: percentage of specific killing= 100%×(RLU of wells without antibody - RLU of detection wells)/RLU of wells without antibody. The data were used to calculate the EC_50_ value using a four-parameter nonlinear fitting model with GraphPad Prism software.

### Granzyme B and perforin examinations

T cells were activated by co-culturing with anti-CD3×CD28 activator beads (2 µl beads added to 1 ml cells) and then expanded. The activating reagent was removed, and the T cells were washed twice with PBS and resuspended in X-VIVO^15^ medium to a density of 2×10^6^ cells/ml, and then added to the wells of the antibody-containing 96-well U-bottom plate at 50 µl per well. Target cells were washed twice with PBS, resuspended in X-VIVO^15^ medium to a density of 2×10^5^ cells/ml, and then added to the wells of the 96-well U-bottom plate at 50 µl per well, with a E:T ratio of 10:1. The YK012 antibody was diluted as follows: first, prepare a starting solution of 4000 pM with X-VIVO^15^, then dilute it in X-VIVO^15^ with a 5-fold gradient continuously, resulting in ten different concentrations. The diluted antibody was added to each well of a 96-well U-bottom cell culture plate at 100 µl per well. Culture medium was added to the eleventh column as negative control well. The plate was then placed in a 37°C, 5% CO_2_ incubator. Cells were harvested and frozen at -80°C for later use at 2, 4, 6, and 24 hours after incubation. After sample collection, Granzyme B was detected using an ELISA kit (Elabscience human GZMB). The samples were diluted 1:3 with culture medium and then 100 µl was added to each well for detection. The plate was read using a BIOTEK Synergy HT reader, with OD450 measurements.

For detection of perforin from activated T cells in the presence of Raji-lp and induced by YK012, the method was exactly same as above-mentioned, except the detection kit was Elabscience human perforin ELISA Kit.

### ADCC and CDC assays

Purified human NK cells were used as effector cells and were mixed with target cells Raji-lp or Jurkat-luc at an E:T ratio of 10:1. They were cultured for 24 hours in the presence of YK012 antibody or anti-CD20 mab (Shanghai Yuanye Bio-Technology Co., Ltd, S28566-1mg) after which fluorescence signal detection was performed. The concentration of YK012 or anti-CD20 mab began at 8 nM, and 4-fold dilutions in ten tiers, plus medium only as zero antibody control. For antibody-dependent cellular cytotoxicity (ADCC) assay, each well contained 100 µl of above antibody YK012 or anti-CD20 mab, 50 µl of effector cells, and 50 µl of target cells. After co-culturing for 24 hours at 37°C with 5% CO_2_ in a cell culture incubator, sixty µl of cells were transferred to a white plate, and 60 µl of luciferase substrate was added to detect the fluorescence signal (RLU). An anti-human CD20 antibody (IgG1) was used as a positive control.

The Raji-lp target cells were seeded at a density of 1×10^4^ cells/50 µl into a 96-well U-bottom cell culture plate. YK012 and anti-CD20 antibody (IgG1), started from 80 µg/ml were diluted by 5-fold over 10 gradient concentrations. Then, 100 µl of each diluted antibody was added to the cell wells. The cells were incubated at 37°C with 5% CO_2_ in a cell culture incubator for 30 minutes to allow the antibodies to maximally bind to the cells. For complement-dependent cytotoxicity (CDC) assay, 50 µl of human non-lysed serum was added to each well, mixed gently and incubated for 4 hours. The cells were then suspended evenly by aspiration to take 60 µl to a non-transparent white 96-well plate. 60 µl of luciferase substrate was added, and the fluorescence signal value (RLU) was measured using the Fluorescence mode of the BIOTEK Synergy HT reader.

### YK012 *in vivo* efficacy study in CIA model

On Day 0 and Day 21, 200 μL of an emulsion containing 0.2 mg of type II collagen mixed with adjuvant was injected intradermally at the base of the tail of the hCD3EDG/hCD19 transgenic mice. A total of 33 male hCD3EDG/hCD19 transgenic mice aged 8 weeks were included in the study, with 30 mice assigned to the experimental groups. Based on the flow cytometry results and CRP levels obtained on Day 26, the mice were divided into five groups of six animals each: a normal control group (Group A), a model group (Group B), and three dose groups (Groups C, D, and E) receiving YK012 at 0.125 mg/kg, 0.25 mg/kg, and 0.5 mg/kg, respectively. Except for Group A, the remaining groups received an emulsion of type II collagen and adjuvant via intradermal injection at the tail root on Day 0 and Day 21 to induce arthritis. Drug administration began on Day 27 in the dose groups and continued until Day 73, with the dosing frequency adjusted during this period for a total of 15 administrations. The pharmacodynamic effect and pharmacological action of YK012 were evaluated through joint clinical scoring, protein level detection, joint pathological staining scoring, and flow cytometry. The animal husbandry and management were conducted in accordance with the SOPs of Suzhou Hekai Biotechnology Co., Ltd., with reference to the Guide for the Care and Use of Laboratory Animals, 8th Edition (2010), and the Animal Welfare Act (Public Law 99-198). The animal use protocol for this study was approved by the Institutional Animal Care and Use Committee (IACUC) of our institution. The IACUC approval number is: A2023003.

### Pharmacological study of YK012 in huHSC-NCG-M mice

HuHSC-NCG-M mice were developed by Gempharmatech. Standard SPF-grade irradiated sterilized genetically modified mouse feed is used. Mice were randomly grouped (6 mice/group) based on body weight and the level of hCD45 × hCD3. The day of grouping is defined as Day 0 (D0). Dosing was performed on D0, D3, D7, D10, D14, D17, and D21. Dosing volume: 10 μL/g × mouse body weight (g). YK012 was given by two different doses, G1 (0.1 mg/kg, mpk) or G2 (0.5 mpk). Mice were weighed weekly, and GvHD was observed twice. After grouping, all mice were fed nutritional gel daily. Peripheral blood was collected for flow cytometry on D-5, D1, D7 (before dosing), D11, D14 (before dosing), and D21 (before dosing). Detection markers: L/D, mCD45, hCD45, hCD79b, hTCR-α/β, hCD38, hCD138. The endpoint was conducted on D22. During the experiment, if the mice reach the humane endpoints, euthanasia will be carried out in accordance with animal welfare standards, and death will be achieved through inhalation of an excess of 95% CO2.

*Ad libitum* feeding and water-drinking. Each batch of feed is provided with a feed quality certificate by the production unit, and a third-party test report is provided annually; the testing standards refer to the national standards GB 14924.3 -2010 “Nutrient Requirements for Laboratory Animals” and GB/T 14924.2 -2001 “Hygiene Standards for Laboratory Animal Feed”. Drinking water is tested once a month, conducted in -house, and an annual report from the local water company is provided, referring to the national standard GB5749-2022 “Hygienic Standards for Drinking Water”.

B cell counts and percentage data are expressed as Mean ± SD and analyzed using the Log-rank test; other experimental results are expressed as the mean ± standard error of the mean (Mean ± SEM). Data were analyzed using SPSS 24, and comparisons between two groups were made using the independent samples T-test. A P-value of less than 0.05 is considered to indicate a statistically significant difference. Graphing was performed using GraphPad Prism 9.

## Results

### Design, production and binding characterization of the YK012

To develop a novel anti-CD3xCD19 bispecific TCEs, considering the whole spectrum of properties needed for engagers to overcome the challenges during the preclinical, clinical, and marketing phases, we optimized a new platform, FIST to generate symmetrical tetravalent bispecific antibody by fusion the scFv of anti-CD3 with a full anti-human CD19 IgG4 at the C-terminus of Fc region, as depicted in Figure 1, A. The designed molecule YK012 possesses two antigen binding sites for CD19 at the N-terminal, two binding sites for CD3 at the C-terminal, binding to CD19 and CD3 in 2+2 mode, different from many asymmetrical engagers that utilize Knob-into-hole^16^ (KIH) platform or other alike 1+1 platforms. FIST molecules, such as YK012, bear the unique advantages of high expression, easy downstream processing, and robust CMC as classical IgG, significant low cost of manufacturing. YK012 was stably produced in CHO cells and purified by optimal procedures. Quality and stability examinations were performed to assure its qualifications in following studies.

**Figure 1.**
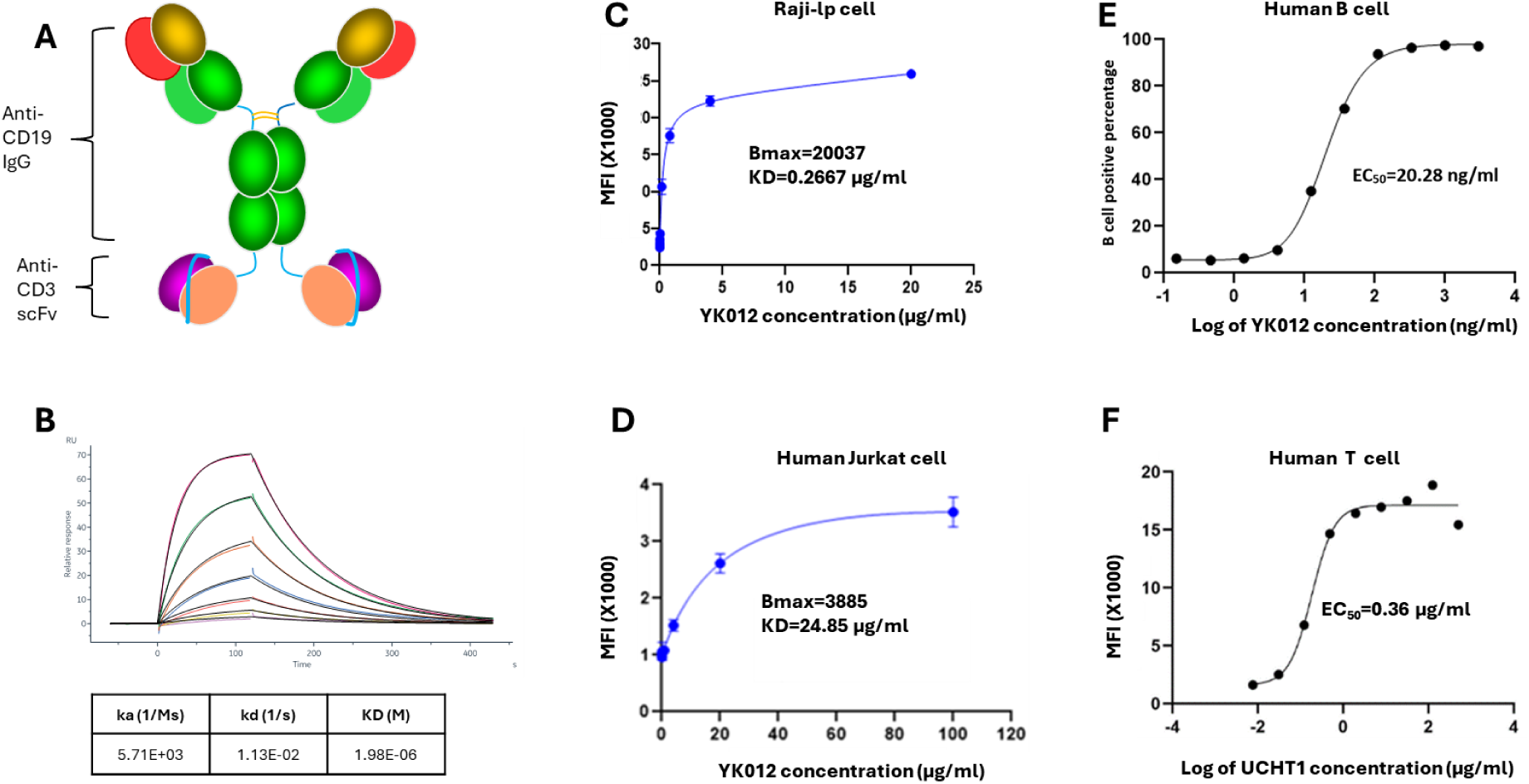
Schematic representation and characterization of YK012. (A) Schematic illustration of the bi-specific anti-CD3xCD19 T cell engager YK012, which contains two scFv of anti-CD3 fused with the anti-CD19 IgG4 at the C-terminal. (B) BIAcore measurement of YK012 to human CD3 ε/δ heterodimer, RU-time curves, and KD. (C) Receptor saturation binding assay of apparent affinity for YK012 to Raji-lp cells. (D). Receptor saturation binding assay of apparent affinity for YK012 to human Jurkat cells. (E) Titrated YK012 binding to human B cells, positive rate verse antibody concentration. (F) Titrated UCHT1binding to human T cells, MFI verse concentration.

By placing the CD3-binding scFv at the C-terminus of Fc, its binding ability to CD3 target was intentionally weakened to lower the stimulation potency to T cells in the “B-YK012-T” synapses. The intrinsic affinity (KD) of YK012 to CD3 was measured by BIAcore. YK012 was captured by CM5 chip, varied concentration of CD3 ε/δ heterodimer as the flowing phase. Response unit (RU) curves of different concentrations were recorded (Figure 1, B). Through 1:1 mode of interaction, the intrinsic KD was calculated to be 1.96×10^-6^ M. The low KD of YK012 to CD3 was further examined by cell-based methods. The value is way below the apparent KD of the parental clone, UCHT1 (Figure 1, F), which was roughly 1.8 nM (0.36 µg/ml). C-terminal fusion of UCHT1 scFv has been reported to have lower KD compared to N-terminus fusion^17^. In receptor saturation assay with Jurkat cells, the apparent KD of YK012 to CD3 in the TCR complex was 24.85 µg/ml (Figure 1, D), or around 124 nM. Overall, the cellular binding of YK012 to TCR on T cells was confirmed to be moderate.

Next, we used flow cytometry to determine the binding activities of YK012 to CD19 in Raji-lp cells (Figure 1, C) and human B cells (Figure 1, E). HD37 mAb was reported to have an intrinsic KD of 4.1±0.3 nM to CD19 antigen^18^. There should be no detrimental impact of CD19 binding strength when the same antibody sequence was adopted as IgG4 in YK012. In fact, the KD measured by BIAcore for YK012 to human CD19 is about 0.5 nM (data not shown). At cellular level, the apparent affinity by means of receptor mean fluorescence intensity (MFI) of YK012 to CD19 on Raji cells was 0.2667 µg/ml, or 1.3 nM. When using titrated YK012 on human B cells with corresponding positive rate, the EC_50_ was 20.28 ng/ml. Nevertheless, the difference of cellular binding apparent affinities of YK012 to membrane CD19 and membrane CD3 reached ∼100-fold (1.3 nM vs. 124 nM), allowing sequential rather than simultaneous binding of YK012 to the two types of cells in blood stream. Meanwhile, the low binding strength of YK012 to TCR means if any, only unstable antibody-antigen complex could form should single YK012 molecule encounter a T cell.

### YK012 activates T cells requiring the presence of CD19^+^ target cells in a dose-dependent manner

The essential function of any engager is to crosslink immune cells with target cells to exert immune mobilization and redirection. To confirm such capacity of YK012, we investigated its feature to activate T cells in the absence or presence of CD19^+^ target cells, such as Raji. PBMC derived T cells, once activated, express increased amounts of biomarkers, such as the early marker CD69 and the late marker CD25. While both CD4^+^ and CD8^+^ T cells express CD25, the latter belongs to cytotoxic T cells, which are the main sources of cytotoxicity.

As the data showed, using CD25 as a marker to check T cell activation in FACS, it was evident that without the presence of CD19^+^ target cells, only YK012 and T cell incubation (YK012 + T) could barely activate CD8^+^ T cells. The CD25^+^CD8^+^ subset accounted for 0.3% (Figure 2, A, C). In contrast, with the addition of CD19^+^ Raji cells, i.e., the incubation of three components (YK012 + T + Raji), YK012 induced significant CD25 expression. The CD25^+^CD8^+^ subset now accounted for 10.9% (Figure 2, B and C). Upon induction of YK012 to T cells, CD25 expression in other subset populations (CD8^-^, which included CD4^+^) also drastically increased.

**Figure 2.**
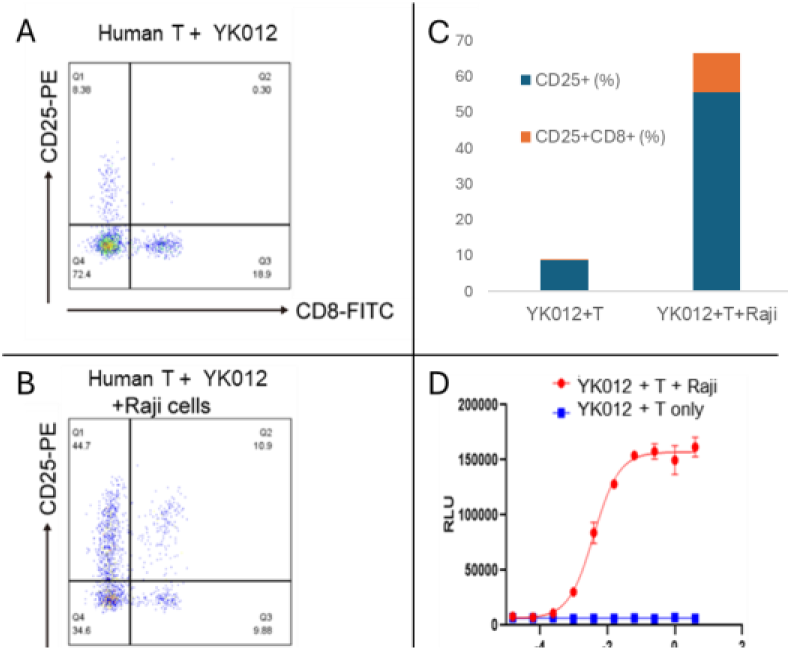
CD3xCD19 activates T cells and requires the presence of CD19^+^ Raji cells. Face analysis of the surface expression of CD25 on CD8^+^ T cells among PBMC derived human T cells stimulated by YK012 without (YK012+T, **A**) or with (YK012+T+Raji, **B**) the presence of Raji cells. The increase of CD25^+^ percentage in the two settings was shown (**C**). YK012 stimulation of T cell activation was dose- and Raji-dependent (**D**).

To find out if there was a dose-dependent correlation between T cell activation and YK012 concentration, a NFAT-responsive reporter cell line, E 6.1-NFAT-luc derived from human T cell Jurkat, was constructed by expressing the luciferase under the control of NFAT promoter. In T cells, NFAT proteins are activated following TCR ligation. In the presence of CD19-expressing cells Raji, addition of YK012 formed bundles through CD19 binding and crosslinked TCR in Jurkat E 6.1-NFAT-luc cells, resulted in the activation of TCR complex as well as subsequent downstream NFAT pathway, which promoted the expression of luciferase under NFAT promoter. As shown in Figure 7.2, D (red line), the reporter signal intensity correlated with dose increment of YK012. In contrast, in the absence of target cells, even when the concentration of YK012 spanned more than 1000 times, no change in fluorescence signal was detected (Figure 2, D, blue line), implying that free YK012 molecule, even as bivalent format, did not activate the NFAT pathway in T cells.

### T cells in PBMC underwent proliferation induced by YK012

Once activated, the T cell undergoes clonal expansion, which is a rapid division and proliferation, which can be seen by the serial reduction of CSFE-derived fluorescence intensity^19^. As cells undergo division and proliferation, the fluorescent dye is evenly distributed to the daughter cells in the second generation. Consequently, the fluorescence intensity of these cells is halved compared to the parent generation. This pattern continues with cell division, the fluorescence intensity of the third generation of cells being further halved compared to the second generation.

CFSE-labeled human PBMC cells (which contained B and T cells) were stimulated by YK012 in the presence of target cells. As control, panT without B cells were also stimulated with YK012 accordingly. The experimental results were presented in Figure 3. The positive control group (Figure 3, C and D, blue peak) showed a synchronous cell division, with the fluorescence values of the dividing cells skewed towards the left for both CD4^+^ and CD8^+^ subpopulations. The negative control group, panT without the presence of B cells (Figure 3, E and F, red peaks) had a main peak at the most fluorescent right, with the highest fluorescence value and only one peak, indicating that panT did not undergo division and proliferation in the absence of B cells.

**Figure 3.**
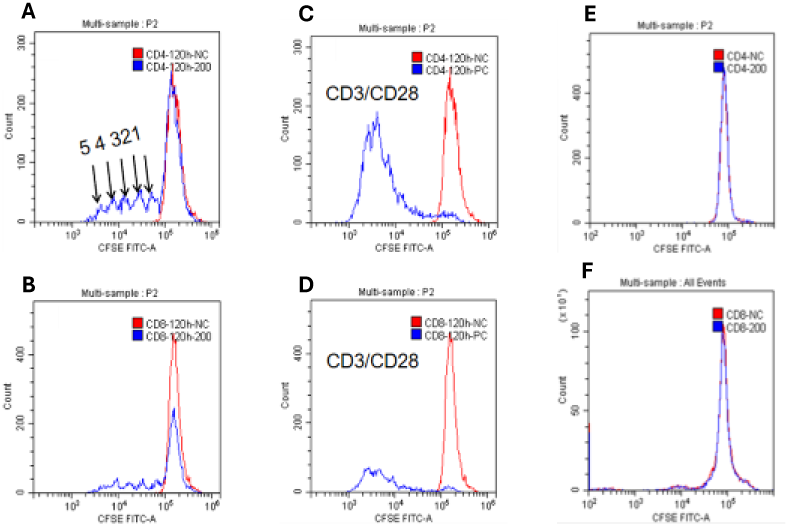
PBMC underwent proliferation induced by YK012. FACS analysis of PBMC or panT induced by YK012. **A** and **B**. CD4^+^ or CD8^+^ subset in PBMC proliferated average 5 times after 120 hours. Blue peaks indicated generations of proliferation. **C** and **D**. CD4^+^ or CD8^+^ subset in PBMC stimulated by anti-CD3×CD28 beads. Synchronized divisions were observed. **E** and **F**. panT subset CD4^+^ or CD8^+^ did not proliferate when incubated with YK012.

For PBMC, it clearly indicated that when YK012 was added, after incubation for 120 hours, PBMC cells underwent division and proliferation, both in CD4 and CD8 subsets. Around 5 generations of division occurred during the 120 hours period. The PBMC cells produced division peaks with decreasing fluorescence intensity (blue peaks in Figure 3, A, B). Across the concentrations of YK012 tested, the number of divisions in PBMC did not show a dose-dependent manner (data not shown).

### YK012 induced less cytokine release *in vitro* than Blincyto

Cytokine release storm (CRS) is a typical adverse event that most T cell engagers encounter in human applications^20^. Grade 3 or above CRS requires immediate attention from medical professionals to avoid irreversible outcomes^21^. Attenuating T cell stimulation, as applied in current molecule YK012, is one of the main strategies to reduce the frequency and intensity of CRS. Meanwhile, prophylactic pretreatment of patients with immune suppressors, such as dexamethasone, could have additive effect to further alleviate clinical symptoms of CRS. YK012 was designed to be milder than Blincyto^®^ in stimulation of immune system.

To measure and compare the cytokine level release of activated PBMC cells conjugated by YK012 or blinatumomab with Raji cells, with and without dexamethasone, supernatant from the incubated cell mixture of human PBMC and Raji cells, in the presence of diluted YK012 or Blinatumomab, was collected and measured by an ELISA method. The major cytokines, including IL-2, TNF-α, IFN-γ, IL-6, were individually quantitated and plotted against YK012 antibody concentrations (Figure, 4).

**Figure 4.**
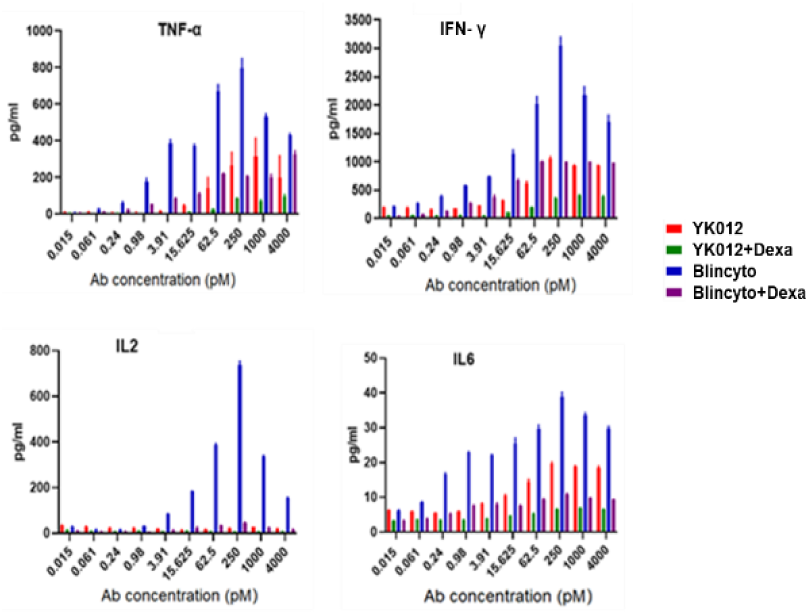
CRS released by PBMC mediated by YK012 or Blincyto^®^ biosimilar, with or without dexamethasone treatment. CD19^+^ Raji cells were co-cultured with activated human PBMC cells at an E:T ratio of 10:1 in the presence of varying concentrations of YK012 or Blincyto^®^ biosimilar, with or without dexamethasone, for 48 hours. Cytokines of TNF-α, IFN-γ, IL-2, IL-6 were individually quantitated by ELISA method and plotted against antibody concentration.

The cytokine release data in the study highlighted that while the levels of cytokines induced in YK012 groups (figure 4, red bars) were generally significantly lower than those of Blincyto^®^ biosimilar (figure 4, blue bars), the application of dexamethasone demonstrated huge reduction in both situations across the wide range of antibody concentrations (figure 4, green verse violet bars). This implies that the future mitigation of clinical severe CRS risk by the preventive prescription of dexamethasone for the usage of YK012 to treat B-cell related abnormalities.

### YK012 mediated T cell killing of CD19^+^ cells over time and not antigen-density dependent

After crosslinking of effector T cells with target cells mediated by T cell engagers, it initiates cascade of reactions from T cell activation, cytokine release to lysis of anchored target cells. How long the synapse may last and how complete the target cells be killed may be dependent on various factors. In the research, incubation time of T cells with different target cells (Raji-luc or NALM6-luc) in the presence of a series of diluted YK012 was investigated to examine the EC_50_ of YK012-mediated killing.

Two cell lines with different antigen densities, Raji-luc (high CD19 expression) or NALM6-luc (low CD19 expression) together with serial diluted YK012 and activated human T cells were incubated for varied length of incubation time (2, 4, 6, and 24 hours) under a fixed E: T ratio of 10:1 to obtain specific lysis percentages. The lysis percentages of YK012-mediated T cells against Raji-luc at different incubation times and concentrations are shown in the table below each figure (figure 5, left); the lysis percentages of YK012-mediated T cells against NALM6-luc at different incubation times and concentrations are shown in the table below each figure (figure 5, right).

**Figure 5.**
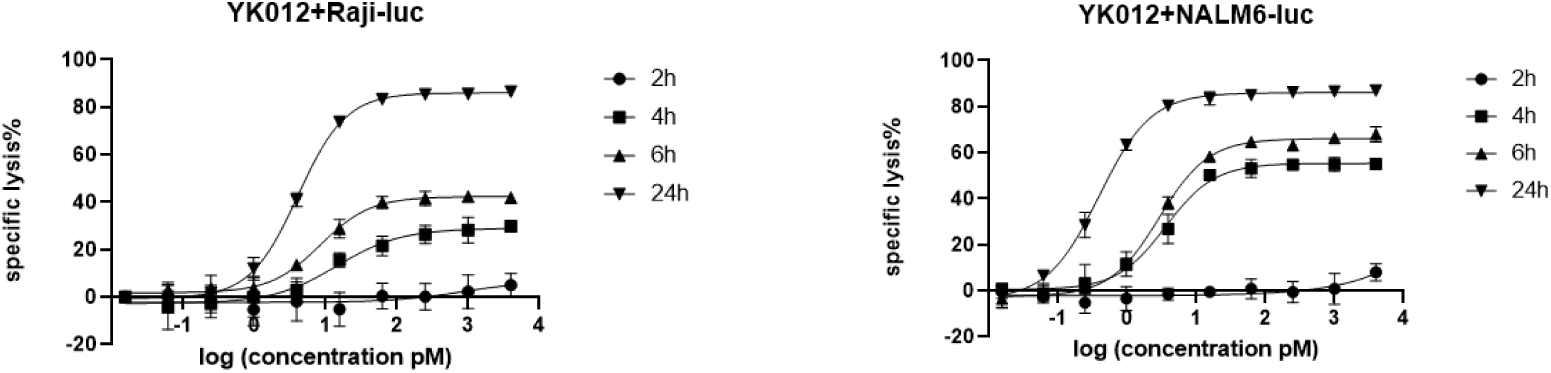
Killing efficiency of YK012-mediated T cell cytotoxicity. Killing efficiency (lysis percentage) of cytotoxicity mediated by varied YK012 concentrations on different CD19-expressing target cells and different incubation times, target cells of Raji-luc (above, left) and target cells of NALM6-luc (above, right).

Time matters in terms of T cell cytotoxicity. The lysis percentage of YK012-mediated killing was very low within the 2-hour incubation period, and even a very high concentration was not helpful (figure 5, black circle curves). With incubation time extended, the killing efficiency gradually increased, and the dose-killing effect relationship gradually became evident. NALM6-luc seemed to be more susceptible than Raji-luc, implying antigen density does matter in affecting YK012 potency. Overall, at a YK012 concentration of more than 10 pM, even a very low mediator molecule concentration can lead to the almost complete killing after 24 hours of incubation (figure 5, upside down black triangle curves). High CD19 antigen density may have negligible beneficial efficient killing mediated by YK012.

### YK012 mediated T cell expression of Granzyme B and perforin

Accompanying the activation of T cells mediated by YK012, toxic molecules are produced as “magic bullets” to exert killing functions. The pattern and potency of YK012 mediated release of toxic molecules, represented by perforin 1 and granzyme B, were investigated *in vitro*. When human T cells and Raji-lp cells were mixed and cultured with various concentrations of YK012, the supernatants collected at different time points were detected for the expression and release of granzyme B by ELISA. Results (Figure 6, left) showed the level of granzyme B began to increase as early as 2 hours after culture, and increased over time, lasting up to 24 hours. Dose-dependence up to 62.5 pM and time-dependence up to 24 hours of granzyme B expression induced by YK012 were prominent (Figure 6). For concentrations of YK012 over 62.5 pM, the amount of granzyme B started to drop, implying that there is a range for the dose-dependent release.

**Figure 6.**
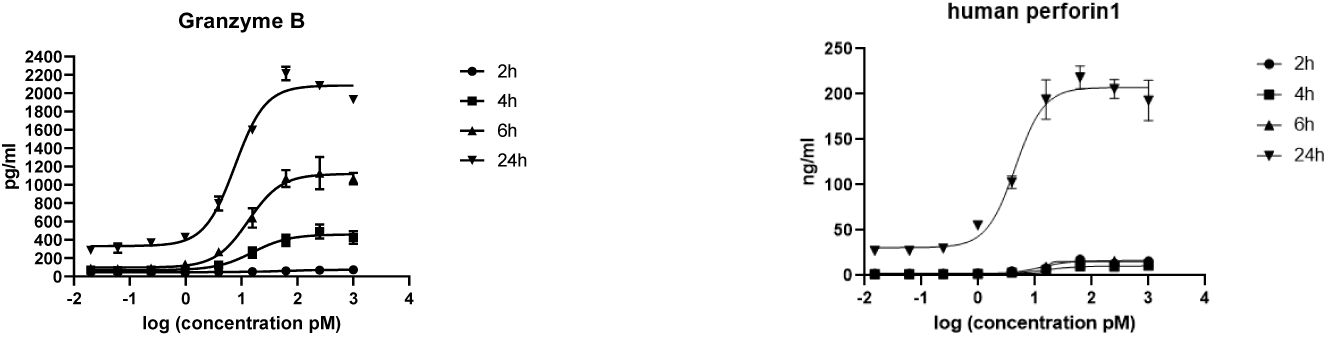
YK012 mediated T cells release of Granzyme B and perforin. In the presence of Raji-lp cells, T cell release of granzyme B (left) and perforin 1(right) at different incubation periods (2,4,6 and 24 hours) mediated by varied YK012 concentrations.

The other important toxic molecules of activated T cells often release is perforins. The pattern and potency of YK012 mediated release of perforin 1 was investigated *in vitro*. In the experiments, after treatment with YK012, supernatants from samples collected at various times were tested for perforin 1 expression and release using the ELISA method. Perforin 1 expression was detectable as early as 2 hours post-treatment at an appropriate YK012 concentration, however, the expression level was generally very low shorter than 24 hours post-treatment. At the 24 hours timepoint, there was a clear dose-dependent relationship and significant expression of perforin 1(Figure 6, right). This study demonstrated that in the presence of CD19+ target cells Raji-lp, YK012-treated T cells exhibited time and dose-dependent perforin expression; release of perforin seemed to be the late event of T cell activation.

### YK012 has minimal ADCC and CDC functions

The YK012 molecule contains the IgG4 Fc domain, which has moderate binding affinity for Fcγ receptors (data not shown). ADCC is a specific killing mechanism where NK cells bind and kill IgG-coated target cells through their Fcγ receptors. To assess whether YK012 would directly mediate NK cells, without the help of T cells, to kill cells through Fc – Fcγ receptors, NK cells were co-cultured with Raji-lp and Jurkat-lp cells, respectively in the presence of varied concentrations of YK012. The positive control was a CD20 (IgG1) antibody. Results are shown in Figure 7 (left). The assay demonstrated that the CD20 monoclonal antibody (IgG1 type), which also recognizes Raji-lp cells, could mediate NK cell killing of Raji-lp efficiently through ADCC (Figure 7, left, black dot curve), while YK012 exhibited a little bit of killing only at the highest concentration (4 nM, Figure 7, black square curve). YK012 did not mediate NK cell killing of CD3-positive Jurkat-lp cells, even at the highest concentration (Figure 7, left, black triangle curve), indicating that YK012 would mediate minimal ADCC killing of human T cells at normal physiological conditions.

**Figure 7.**
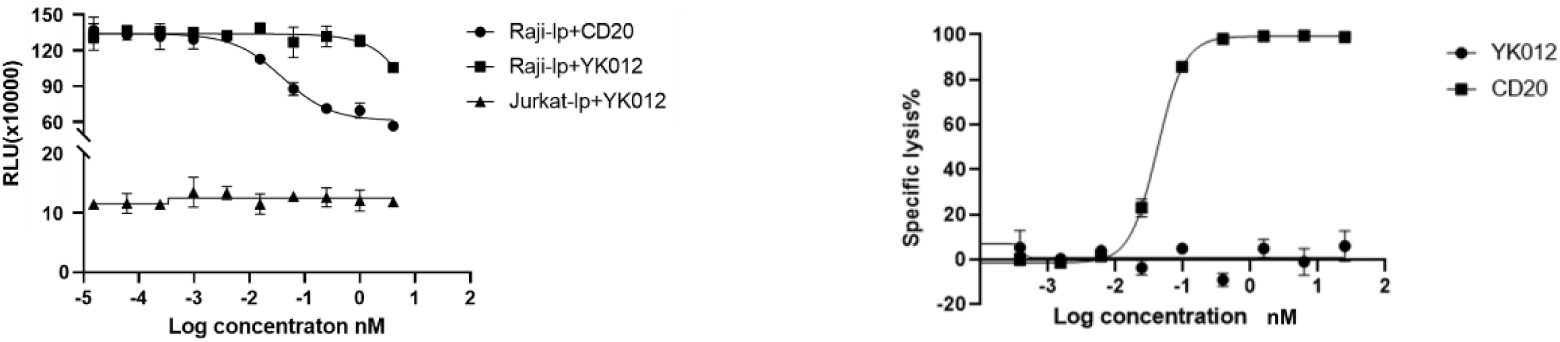
ADCC (left) and CDC (right) function examination of YK012. In the presence of NK cells, varied concentration of YK012 incubated with Raji-lp (left, black square) or Jurkat-lp (left, triangle), respectively, for measurement of ADCC activity or CDC activity. Anti-CD20 IgG1 was included as positive control in each assay.

The YK012 molecule contains the Fc structure of IgG4, which can bind to complement C1q. According to literature reports, the affinity between IgG4 antibody monomers and C1q is very low, approximately 100 µM^22^. During the interaction of YK012 with CD19^+^ and/or CD3^+^ cells, the antibody-antigen complex can potentially induce complement-dependent cytotoxicity (CDC). To investigate whether the Fc segment of YK012 can mediate CDC effect in causing the lysis of target cells, Raji-lp cells (CD19 positive target cells) were incubated with different concentrations of YK012 and human serum (not inactivated), and the lysis percentage of Raji-lp cells was measured. Since Raji-lp cells also express CD20, a CD20 antibody (IgG1) was used as a positive control for CDC assay. The results are shown in the figure below (Figure 7, right, black square).

As shown in the figure 7 (right plot), the CDC effect mediated by the CD20 antibody was very significant, with Raji-lp exhibiting the typical antibody concentration-dependent lysis effect (S-shaped curve, black squares). In sharp contrast, over a wide range of concentrations, YK012 did not show dose-dependent curve or that of the CD20 antibody, and Raji-lp (expressing CD19) did not show any detectable lysis (Figure 7, right, black circles,), proving that the Fc part of YK012 does not mediate CDC effects against target cells. Since the CD19 binding unit of YK012 has a significantly higher affinity for B cells than its CD3 unit has for T cells, it can be reasoned with certainty that YK012 may only mediate negligible CDC effect against human T cells.

### YK012 alleviated the symptoms of CIA mice *in vivo*

The CIA model study was entrusted to Suzhou Hekai. The chicken type II collagen-induced mouse arthritis model is a commonly used model for studying human arthritis diseases and serves as an important tool for investigating the pathogenesis of arthritis. CD3 and CD19 are crucial targets in the treatment of arthritis diseases. Since the test substance is a humanized antibody with no cross-reactivity with mice, hCD3EDG/hCD19 transgenic mice on a C57BL/6 background were selected for model induction. There are no alternative non-live animal tests that can replace this experiment.

During the experiment, the body weights of the mice in all groups fluctuated within the normal range (data not shown). Joint clinical scoring revealed a downward trend in the joint scores of the mice in the dose groups compared to the model group (Figure 8), but no significant differences were observed. Protein level detection did not reveal significant differences in CRP (C-reactive protein), RF (rheumatoid factor), ACPA (anti-cyclic citrullinated peptide antibodies), and SAA (serum amyloid A) levels among the groups (data not shown). Joint pathological staining scores also showed a downward trend but without statistical significance. Flow cytometry results demonstrated changes (fluctuations) in the absolute counts of peripheral blood B cells (CD20+/B220+) and T cells (TCRβ+) as well as their proportions, beginning on Day 33 (6 days after the first administration) and partially or fully recovering in the later stages of the experiment, but statistically not significant (data not shown).

**Figure 8.**
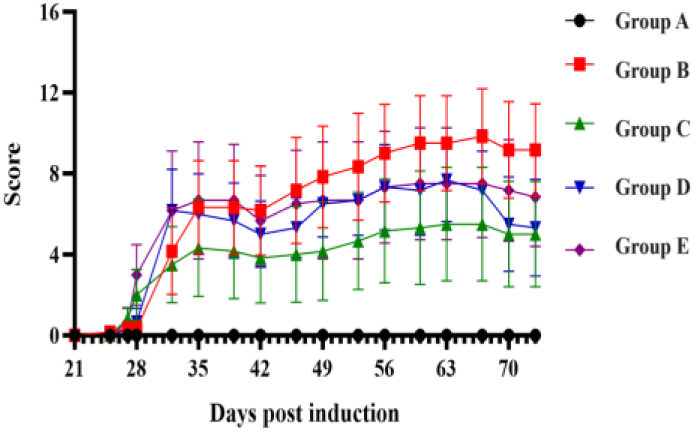
Clinical joint score of CIA mice treated by YK012. Group A (black dot) and B (red square) are controls of normal mice (no collagen, wild type) and CIA mice without YK012 treatment, respectively. Group C (green up triangle, 0.125 mpk), group D (blue down triangle, 0.25 mpk) and group E (cyan diamond, 0.5 mpk) are YK012-treated groups at different doses.

### Significant B cell depletion *in vivo* in HuHSC-NCG-M mice treated by YK012

The important secondary lymphoid organ, the spleen, is better reconstructed in HuHSC - NCG-M mice, and the volume and weight (organ-to-body weight ratio) of the spleen are significantly increased compared to HuHSC-NCG mice. Moreover, the proportion of hCD3+ T cells in the spleen is increased, which is closer to the proportion of T cells in the human spleen. In this situation, the anti-drug antibody response against YK012 would be significantly reduced after repeated usage, allowing the accumulation of B cell depletion to occur.

Right before grouping, the average level of hCD45 × hCD3 in mice was 3.31%. During the study, the body weight of mice in different groups almost unchanged (body data not shown). After administering YK012 via tail vein injection twice a week for a total of 7 doses, there was a significant reduction in the number of B cells (CD79b+) in the peripheral blood of huHSC-NCG-M mice. The B cell depletion effect of the G1 (0.1 mpk dose) group was superior to that of the G2 (0.5 mpk dose) group (Figure 9). No mice were euthanized during the experiment due to the welfare issues, nor unexpectedly death during the experiment.

**Figure 9.**
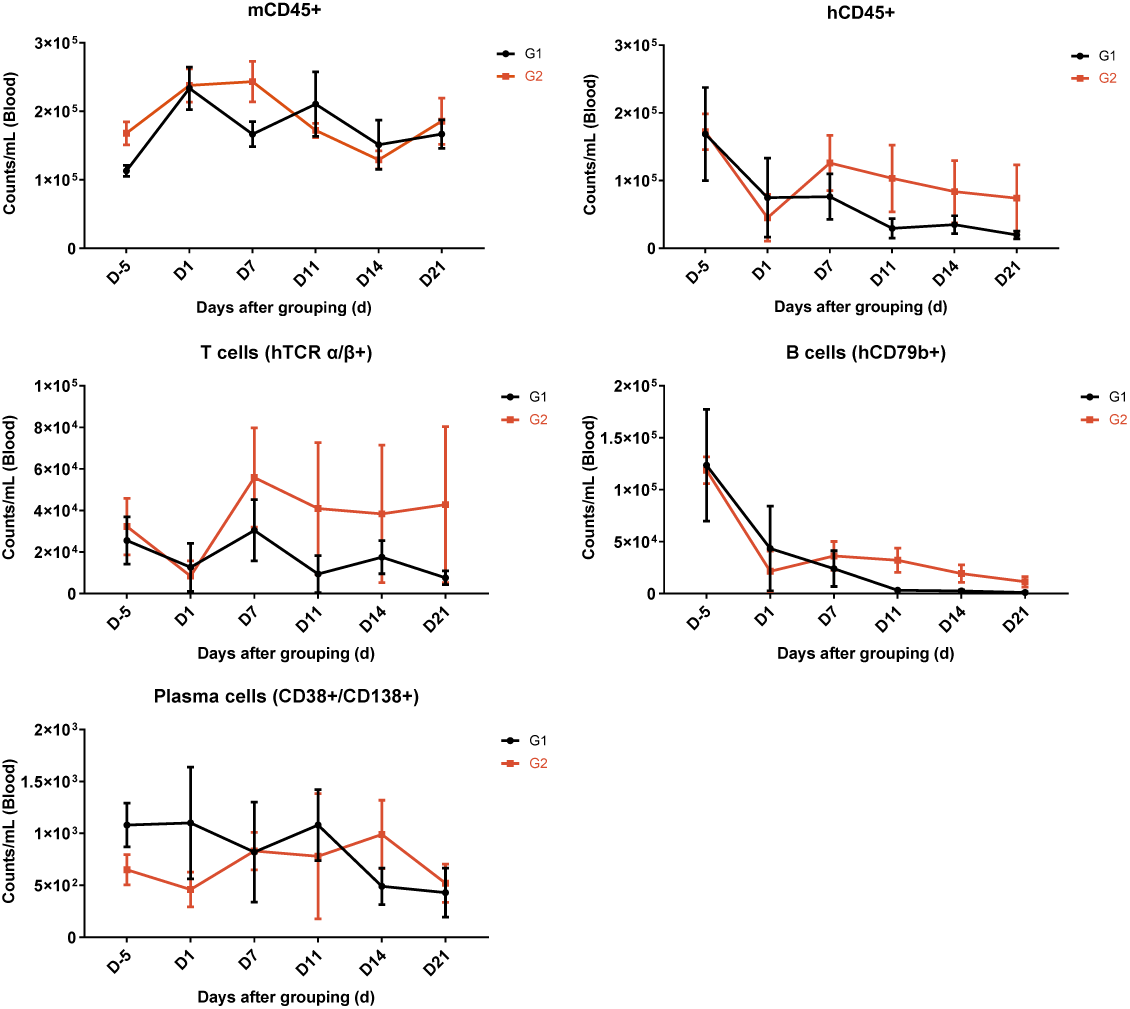
Pharmacological study of YK012 in huHSC-NCG-M mice. Changes of the counts of immune cells (mCD45^+^, hCD45^+^, T, B and plasma cells) in the peripheral blood of huHSC-NCG-M mice treated by high (0.5 mpk, red lines) or low (0.1 mpk, black lines) doses of YK012 over time (day). The data are presented as Mean ± SEM.

The statistical analysis of the peripheral blood flow cytometry results demonstrated that in all YK012 treated groups, compared to D-5, the proportion and number of B cells (CD79b+) on D1, D7, D11, D14, and D21 all showed depletion, and the degree of decrease deepened progressively with the extension of time and the increase in the number of doses administered. In Group G1, compared to D-5, the proportion of B cells (CD79b+) on D7 showed a significant decrease (P<0.05); the proportion of B cells (CD79b+) on D11, D14, and D21 showed a more pronounced significant decrease (P<0.001). In Group G2, compared to D-5, the proportion and number of B cells (CD79b+) on D1 and the number of B cells (CD79b+) on D7 showed a significant decrease (P<0.05); the proportion of B cells (CD79b+) on D7 and the proportion and number of B cells (CD79b+) on D11, D14, and D21 showed an even more pronounced significant decrease (P<0.01). Compared to Group G2, Group G1 had lower proportions and numbers of B cells (CD79b+) on D11, D14, and D21, and showed a significant difference on D11 (P<0.05). The proportion of T cells (TCR α/β+) on D7, D14, and D21 all showed an increase.

## Discussion

Targeting autoreactive B cells by blocking B cell activation and/or depletion of B cells has shown promise for the treatment of multiple autoimmune diseases. However, a large proportion of patients do not respond well to the therapies^3^ or released after initial successes. In this study, we proposed that using a novel tetravalent anti-CD3xCD19 (YK012) could be more efficient for the killing of autoreactive B cells and provide overall benefit for the autoimmune patients compared to monoclonal antibody therapy or CAR-T therapy. In this study, we designed and generated a symmetrical tetravalent TCE targeting CD19 and CD3, YK012, based upon the in-house FIST platform. We confirmed that while the bivalent-CD3 binding YK012 has much reduced intrinsic affinity to CD3 compared to parent antibody UCHT1, it would only activate T cells in the presence of CD19^+^ target cells, promoted the proliferation of PBMC, and effectively elicited T cell-mediated cytotoxicity only to CD19^+^ cells. Meanwhile the *in vitro* released major cytokines associated with CRS were minimized compared to Blincyto^®^ biosimilar, additional reduction was possible when accompanied using dexamethasone. More importantly, YK012 mediates the cytotoxicity of activated T cell to CD19-expressing target cells by releasing toxic molecules such as granzyme B and perforin. In animal studies, YK012 exhibited the ability to alleviate symptoms of CIA mice, induced almost complete depletion of peripheral B cells in huHSC-NCG-M mice. These data demonstrated that the engineered YK012 has the potential to be more effective but less toxic for the treatment of autoimmune diseases by killing CD19^+^ B cells.

Nowadays the approved approach for B cell targeting is using anti-CD20 monoclonal antibody to deplete B cells^23, 24, 25^. Although it is highly efficient to deplete B cells in the periphery, it is less efficient to deplete autoreactive B cells in the lymphoid organs and inflamed tissues^6^. Currently, it is not well understood why the Fc-mediated ADCC of B cells is impaired in those tissues. Autoreactive pre-B cells and plasma cells have been shown to mediate the pathogenesis of autoimmune diseases and manifestations of symptom and autoimmunity^6, 26, 27^. The lack of or low CD20 expression in pre-B cells and plasma cells could be another reason for patients not responding to anti-CD20 treatment^6, 26, 27^. In contrast, CD19 is broadly expressed throughout B cell linage including pre-B cells and plasma cells and acts as a critical regulator for B cell activation and function^11, 12, 28^. This suggests targeting CD19 may have more advantages than targeting CD20. In line with this, a non-depleting antibody LY3541860 potently inhibited B cell function and showed superior activity in the inhibition of disease severity in a TD1 mouse model, compared to anti-CD20^29^.

The TCE targeting CD19 approach has been approved for leukemia. Blinatumomab is the first BiTE approved by the FDA for the treatment B-ALL^15^. It is composed of two scFv connected with a flexible linker with small size (55 kDa), which leads to quick kidney clearance and requires continuous infusion. But it has limitations owing to its extremely short half of slightly over 2 hours and strong adverse events including CRS and neurotoxicity. The short space between anti-CD3 scFv and anti-CD19 scFV was attributed to its strong activation ability on T cells^30^. The distance between the two antigen binding sites has been proposed to be a strength regulator for the anti-CD3 arm to induce storm-like T cell activation^30^. Here, we engineered a TCE that contains a full anti-CD19 IgG4 with fusion of anti-CD3 scFV at the C-terminal. Compared to the parent anti-CD3, the molecule we designed has less binding affinity to CD3 as well as longer distance between the two binding sites, suggesting that it may induce weak T cell activation and has less side effect. Further, the engineered YK012 only elicited killing on CD19^+^ Raji B cells but not CD19^-^MV-4-11. This suggest YK012 would not induce non-specific killing, thus further limits its potential systematic toxicity.

Immunotherapy modalities that harness the cytolytic potential of T cells to target and kill tumor cells, such as Chimeric antigen receptors (CAR) -T cells or BiTE, have been intensively studied for the treatment of a variety of cancers^14, 31^. Only recently have CAR-T cells been investigated for autoimmune diseases. In 2021, a woman with severe systemic lupus erythematosus was treated with CD19-targeting CAR-T cells^32^. The infusion of CAR-T cells significantly reduced the disease severity and showed acceptable safety^32^. These data demonstrate it is feasible to harness T cells to kill autoreactive B cells in clinic. However, preparation of autologous CAR-T cells requires intensive labor and is time consuming. In contrast, bi-specific antibody is convenient to manufacture and can be easily administrated to patients. Currently it has not been reported that using BiTE or alike to harness the T cytotoxicity to kill the autoreactive B cells. Our candidate YK012 has strong potential to further improve the treatment outcomes for autoimmune diseases.

This study has several limitations. While YK012 demonstrated some extent of symptoms relief in autoimmune CIA model, due to anti-drug antibody generation, it was not able to generate accumulated improvement following repeated injection of the TCE. YK012 bears fully human constant region sequence and humanized antibody variable region, likely induced strong immune response, including anti-YK012 antibody production by the murine immune system. In another murine model where, human immune cells were developed, although repeated infusion of YK012 resulted in deepened depletion of peripheral B cells, due to the relatively short life span of this model (< 1 month), depletion of B cells from secondary lymphoid tissues could hardly be achieved, which may be critical in terms of generating beneficial outcomes for autoimmune patients. A fully murine surrogate molecule may overcome the above restrictions. Nevertheless, B-cell depletion starting from peripheral blood is the right start of the therapy, long-lasting of B-depletion likely ends up with improvement of autoreactive symptoms, as shown in CAR-T trials in SLE patients^33^.

One big concern when repositioning TCE from cancer patients to autoimmune patients lies in the safety issue. In the new situation, many factors such as CRS level, lymphocytes composition, T cell status, distribution of autoreactive B cells etc., may have drastically differences for the appropriate clinical management to be developed, let alone the dosing scheme change. As a promising sign, it has demonstrated that many candidate antibody drugs used in cancer patients can be dosed at similar level in autoimmune populations with much less frequency^34^, highlighting the feasibility of using TCEs in autoimmune diseases. In summary, from current report the engineered YK012 has the great potential to provide an effective treatment option for patients with autoimmune diseases.

## Acknowledgement and declaration

Andy Yuan, Likun Zhao, Lili Bai, Xuning Dai, Yang Wang and Qingwu Meng are employees of Excyte Biopharma Ltd (China). Excyte LLC is a subsidiary company of Excyte Biopharma Ltd. Ray Qirui Yuan and Isbella Yang are summer interns in Excyte LLC. The authors thank Dr. Xiangping Yang for helpful discussion and manuscript revision.

## Notes

### Competing Interest Statement

Andy Qingan Yuan and Qingwu Meng are the co-founders of Excyte LLC and Excyte Biopharma Ltd. Likun Zhao, Lili Bai, Yang Wang and Xuning Dai are current employees of Excyte Biopharma Ltd. Ray Qirui Yuan and Isabella Yang were interns of Excyte LLC.

### Summary of Updates

additional work, contributed by company staff, has generated more data and final entity was named to align with clinical development.

